# In-ear-tACS: Auditory Perception and Side Effects depend on Electrode Montage, Frequency, and DC-Offset

**DOI:** 10.1101/2025.10.30.685487

**Authors:** Luca M. Reinema, Arnd Meiser, Niklas Mellerke, Daniela Rodriguez de Castro, Heiko Stecher, Andreas Radeloff, Christoph S. Herrmann

**Author notes:** Shared first authors (Luca M. Reinema, Arnd Meiser). Shared last authors (Andreas Radeloff, Christoph S. Herrmann).

## Abstract

**Background:** Non-invasive transcranial alternating current stimulation (tACS) of the cochlea might be a promising new therapeutic option for patients with chronic tinnitus. However, electric stimulation of small, sensitive target regions, such as the inner ear, can cause adverse side effects (SEs).

**Objective:** To identify stimulation parameters with low side-effect profiles while reliably stimulating the cochlea, thereby improving patient comfort and safety and paving the way for the development of a medical device for treating patients with chronic tinnitus.

**Approach:** Hearing-healthy participants were stimulated with electrodes in the ear canal. Stimulation of the cochlea elicits a soft hearing impression (HI) in participants, indicating successful stimulation of the early auditory pathway. We systematically compare HS and SEs across distinct electrode configurations and stimulation parameters, including stimulation frequency and the presence of a Direct Current (DC)-offset. We record SE occurrence via visual analog scales after stimulation.

**Main Results:** We find that tACS stimulation between 250 Hz and 2000 Hz reliably elicits HIs in participants. SEs are generally low. No occurrence of SEs resulted in participant withdrawal or severe adverse events, with only phosphenes, skin tingling, and a sense of vibration being reported as impactful. The addition of a slight DC-offset increases the occurrence and magnitude of SEs considerably.

**Significance:** Our results demonstrate the feasibility of tACS in non-invasively stimulating the auditory pathway with minimal adverse SE. Stimulation parameters with a low SE profile can be applied in further studies with tinnitus patients.

## 1. Introduction

### 1.1 Mechanisms of tinnitus

Tinnitus affects approximately 10-15% of the global population and can significantly impair quality of life, particularly in its chronic form (Biswas et al., 2022; Hackenberg et al., 2023; Heller, 2003; Jarach et al., 2022; McCormack et al., 2016). For patients with chronic tinnitus, effective treatment options are currently limited to acoustic interventions, such as masking or habituation techniques, or behavioral training, to develop coping strategies that offer symptomatic relief. A definitive therapy remains to be found (Arts et al., 2016; Bovo et al., 2011; Chang & Zeng, 2012; Czornik et al., 2022; Holder et al., 2017; Jastreboff & Jastreboff, 2003; Mazurek et al., 2022; Park et al., 2023; Quaranta et al., 2008; Ruckenstein et al., 2001). A leading hypothesis for the origin of tinnitus is the *Central Gain Theory*, proposing that altered central auditory mechanisms, often following peripheral hearing loss, lead to the perception of phantom sounds (Auerbach et al., 2014; Dauman & Bouscau-Faure, 2005; Langguth & De Ridder, 2013; Noreña, 2011; Zeng, 2013). This hypothesis is supported by the fact that tinnitus typically manifests at frequencies with the most significant hearing loss (Jastreboff & Jastreboff, 2003).

### 1.2 Transcranial electric stimulation for the alteration of tinnitus

The potential of electrically stimulating the peripheral auditory system was initially identified in cochlear implantation (*CI*). Several studies have shown that electrical stimulation via a CI can lead to a significant reduction in tinnitus symptoms (Arts et al., 2016; Bovo et al., 2011; Chang & Zeng, 2012; Holder et al., 2017; Kaltenbach et al., 2005; Quaranta et al., 2008; Ruckenstein et al., 2001). However, for individuals with normal hearing, CIs are considered too invasive as they involve surgical intervention.

tACS of the inner ear can directly stimulate the early parts of the auditory system. The extension of this treatment approach to tACS focuses on applying alternating current in audible frequency ranges, targeting the cochlea directly to fill in missing auditory stimuli and reduce phantom sounds (Arts et al., 2016; Bovo et al., 2011; Chang & Zeng, 2012; Holder et al., 2017; Quaranta et al., 2008; Ruckenstein et al., 2001).

Among the regions of interest for peripheral stimulation, studies demonstrating the effectiveness of tACS stimulation have primarily focused on stimulating the cochlea (Chang & Zeng, 2012; Suh et al., 2022; Zeng et al., 2011; Zeng et al., 2019; Zeng et al., 2019). Efficacy has been demonstrated even with about 1% of the stimulation reaching the cochlea, using a built-in CI measurement system (Zeng et al., 2019). A wide range of stimulation parameters exists, including frequency and electrode placement, but no consensus on the most effective setup has been reached (Arts et al., 2012; Chang & Zeng, 2012; Suh et al., 2022; Zeng et al., 2019).

### 1.2 Adverse SEs of tACS

Despite its various potential uses, transcranial electrical stimulation (tES) carries the risk of inducing adverse SEs. This is mainly due to volume conduction in the head, which spreads electric currents beyond the targeted tissues. SEs include light phenomena (phosphenes), tingling, and dizziness (Zeng et al., 2019), which hinder the usability of tACS for ambulatory tinnitus treatment. As demonstrated by Zeng et al. (2019), the positioning of the return electrode significantly affects current distribution and the likelihood of adverse effects. Previous research indicates that applying tACS to the vestibular system can cause lateral body sway and auricular stimulation (preauricular, mastoid) may induce dizziness due to proximity to the vestibular system (Antal et al., 2008; Arshad et al., 2014; Bjekić et al., 2024; Laakso et al., 2025; Landgren & Cajander, 2021; Raco et al., 2014). Phosphenes often result from retinal stimulation during frontal or periorbital electrode placement (Antal & Paulus, 2013; Schutter & Hortensius, 2010). Tingling or pricking sensations under the electrodes commonly occur during stimulation (Bjekić et al., 2024; Raco et al., 2014) and are associated with higher frequencies and intensities (Elyamany et al., 2021). However, besides a few exceptions (e.g. Chaieb et al., 2014; Zeng et al., 2019), safety aspects of tACS are investigated in the above-mentioned works in the frequency range of ∼1-80 Hz, with the intent to entrain endogenous brain oscillations. Our work focuses on frequencies in the audible range between 250 Hz and 2000 Hz, aiming not at brain oscillations but at frequencies relevant to the functioning of the cochlea. Our study addresses a gap in our understanding of SE profiles for tACS at higher frequencies. Based on previous literature, we focus our study on seven SE: nausea, muscle twitching, vertigo, headache, phosphenes, skin tingling, and mechanical skin vibration (Elyamany et al., 2021; Matsumoto & Ugawa, 2017).

### 1.4 Study aims

This study addresses the gap identified by Zeng et al. (2019) regarding the SE profile of electrical stimulation, focusing on tACS of the periphal parts of auditory processing (Zeng et al., 2019).

In this study, we aim to examine whether the SEs associated with tACS are tolerated at frequencies in the audible range. We explore differences between when a DC-offset is applied and when it is not (RQ 1.1: Do SEs differ between tACS with DC-offset (tacsDC) and without DC-offset (tacsNO)?). We further explore whether the frequency of stimulation (RQ 1.2 Do SE vary depending on the stimulation frequency?) or the choice of electrode montage (RQ 1.3 Do SE differ between electrode montages?) influences the occurrence and profile of these SEs.

Additionally, our work uses a metric from previous studies: electric stimulation of the cochlea has been shown to produce a faint, pitch-related hearing impression that can be used as an indication of electrical activation of the hearing system (Zeng et al., 2019).

This research aims to investigate whether individuals’ HIs during tACS vary depending on the above-mentioned variables (RQ 2.1: Do auditory impressions differ between tACS with and without DC-offset?; RQ 2.2 Do auditory impressions vary depending on the stimulation frequency?; RQ 2.3 Do auditory impressions differ between electrode montages?).

## Main Section

### 2.1 Methods

#### 2.1.1 Participants

The study was approved by the ethics committee of the University of Oldenburg and pre-registered at the *German Register of Clinical Studies* (DRKS, https://drks.de/search/de/trial/DRKS00033120). All participants provided written informed consent in accordance with the Declaration of Helsinki. Eleven participants aged 18-30 were recruited via an advertisement posted on the University of Oldenburg’s online blackboard. Participants were screened and an audiogram verified normal hearing thresholds (below 20 dB at 125-8000 Hz). Additional exclusion criteria included neuropsychological disorders, metal head implants, skin irritations on the head and non-removable piercings.

#### 2.1.2 Stimulation parameters and equipment

To address *Research Question 1*, two configurations were tested (see Figure 1): (1) in-ear stimulation (right ear) in combination with a forehead return electrode (5*5cm rubber electrode, Neuroconn, Ilmenau, Germany). The return electrode was positioned at FPz, according to the international 10-20 system, and attached to the forehead via conductive paste (Ten20, D.O. Weaver, Aurora, CO). (2) In-ear stimulation with an in-ear return electrode (left ear). In line with *Research Question 2*, both configurations were examined under two conditions: first, using tACS without a direct current (DC) component *(=tacsNO)*, and second, using combined tACS with an underlying DC component (+0.4 mA, *=tacsDC*). We used a battery-driven stimulator with an extended frequency range of up to 3000 Hz (DC Stimulator Plus, Neurocare Group AG, Munich, Germany). A gold-plated tip electrode was used as an in-ear electrode. The experiment administered sinusoidal electrical stimuli at four frequencies (250 Hz, 500 Hz, 1000 Hz, 2000 Hz). Stimulation was administered from 0.2 mA to 1.6 mA in the tacsDC condition and from 0.6 mA to 2.0 mA in the tacsNO condition, both in 0.2 mA steps. The sequence of stimulation intensities was randomized and never exceeded 2 mA.

**Figure 1:**
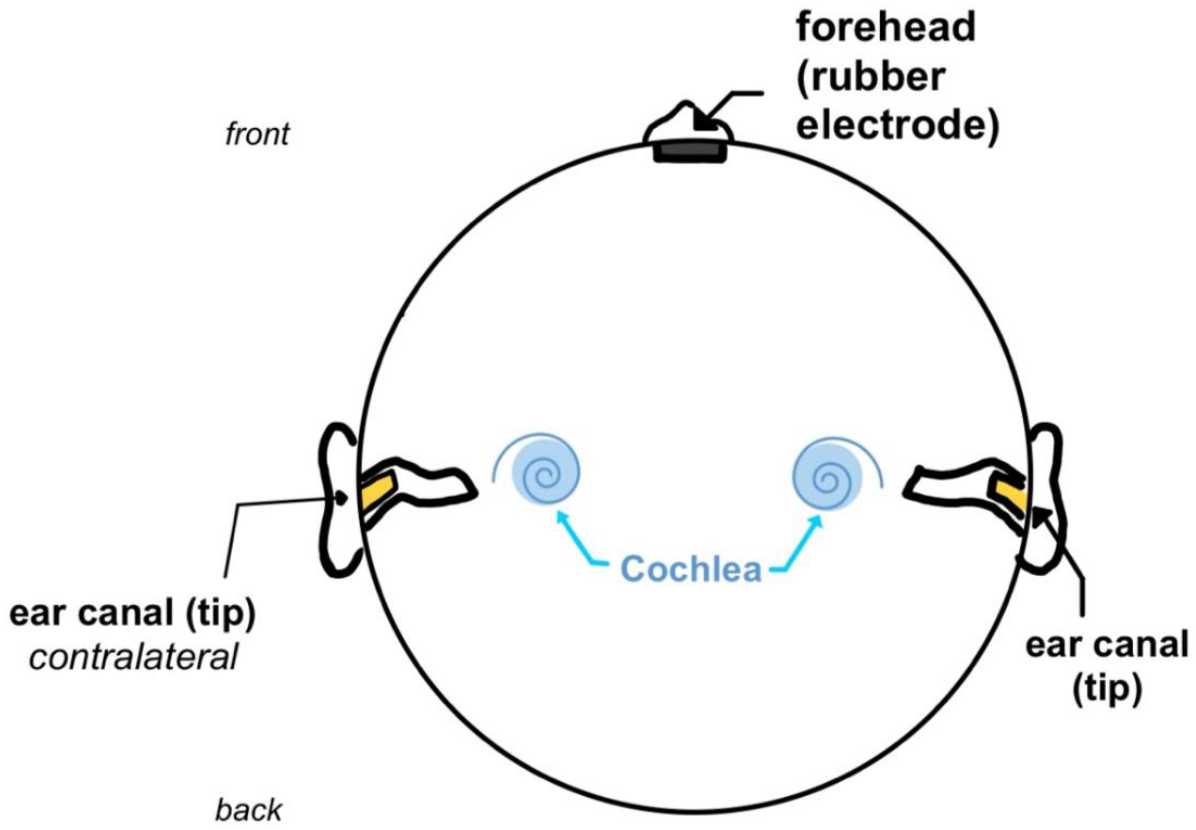
Set-Up - Electrode positions (top view of the head): Stimulation electrode: right ear canal (tip). Return electrodes: forehead (plate), ear canal contralateral (tip), illustration adapted from (Zeng et al., 2019), under a Creative Commons Attribution 4.0 International License, https://creativecommons.org/licenses/by/4.0/.

**Figure 2:**
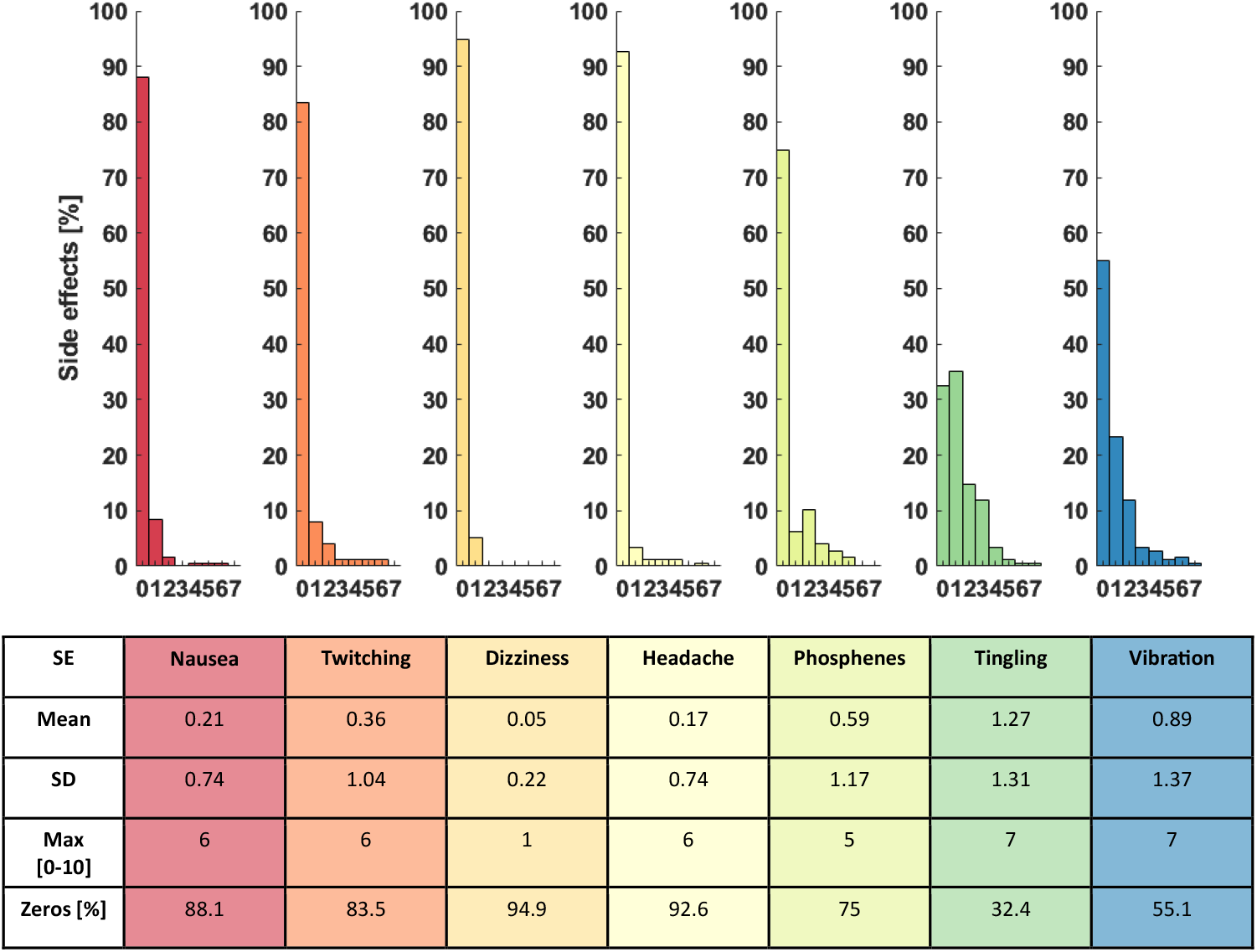
Descriptive Overview: SE scales. Each column shows one SE. On the horizontal axis, we show the rating from 0 to 10, with 7 being the highest observed value. On the vertical axis, we show the percentage of occurrence of each value, so that the values per SE add up to 100%. Below, we show the mean, standard deviation, maximum value, and the percentage of zero ratings, across all participants.

#### 2.1.3 Experimental procedure

Participants were seated in a chair in front of a screen at approximately 80 cm from their eyes. They used a keyboard to start and stop stimulations, advance and abort the experiment and a mouse to indicate SEs. A physician checked the participants’ ear canals for excessive earwax. Impedances of both electrode montages were checked to be below 10 kΩ. Participants completed four training trials in the tacsNO and tacsDC conditions to familiarize themselves with the protocol and the sensations of HI and SE, with a stimulation frequency of 250 Hz.

In line with *Research Question 1*, participants were asked to rate the experienced SE on a Likert-like scale, ranging from 0 (not noticeable) to 1 (slightly noticeable), via 5 (moderately noticeable), to 10 (strongly noticeable). Participants were informed that a 10 rating corresponds to the maximum intensity that was not painful. If any SE became painful, participants were instructed to terminate the session by pressing a button. No participant felt the need to make use of this option. SE scales were presented after all eight randomized intensities within a trial. Participants were instructed to rate their strongest SE sensation during the trial.

In line with *Research Question 2*, after each stimulation intensity, participants were asked to indicate whether they had heard a tone. If they did, they specified whether the tone was heard on the right, left, or on both sides. Finally, participants rated on a scale from 0 (not at all) to 10 (very strongly) how distracted they felt from focusing on the HI. Furthermore, the study used Frenzel glasses whenever participants reported dizziness to objectively detect involuntary eye movements (nystagmus) caused by vestibular stimulation.

The documentation and analysis of the results were conducted using the data analysis software package SPSS (Corp., Released 2023). Figures were produced in MATLAB (R2022b, The MathWorks Inc., Natick, MA, USA).

### 2.2. Results

A total of 11 participants (age range = 21-30 years, *M* = 25.09 years, *SD* = 2.70; seven females) were included in the analysis. A significance level of p <.05 was used for all analyses. We aim to investigate the impact of frequency, DC offset, and position on HI and SE.

Our results show that in-ear tACS reliably elicited auditory sensations with generally mild side effects. The intensity and occurrence of these side effects varied depending on all three variables.

Participants reported (near-) zero values for most SEs, making their practical impact negligible. The analysis focused on parameters with low zero inflation (<80%), such as phosphenes, vibration, and tingling. Given high zero inflation, low mean values (generally below 2/10), and maximum values of 6/10, it can be concluded that the stimulation was generally well tolerated. Descriptive parameters for the SE scales are shown in Figure 1.

We performed an overarching generalized linear mixed model (GLMM) analysis. This approach allows for the consideration of both fixed and random effects, accounts for repeated measures within subjects, and is particularly well-suited to handling non-normally distributed data, as in the case of our SE scales. A Poisson regression with a log-link function was applied to the SE scales. A logistic regression model with a logit link function was used for the binary outcome of auditory perception.

### 1.1 SEs are increased during tacsDC

RQ1.1 addresses the impact of artificial direct current stimulation on adverse SE. Descriptively, the DC offset (RQ 1.1) was always associated with higher ratings for SE (see Figure 3). However, the significance of differences could only be demonstrated for the occurrence of phosphenes (F(1, 214)= 18.88, p<0.001, tacsDC vs. tacsNO B = −2.145, p < 0.001, Exp(B) = 0.12, 95% CI [0.04, 0.31]).

**Figure 3:**
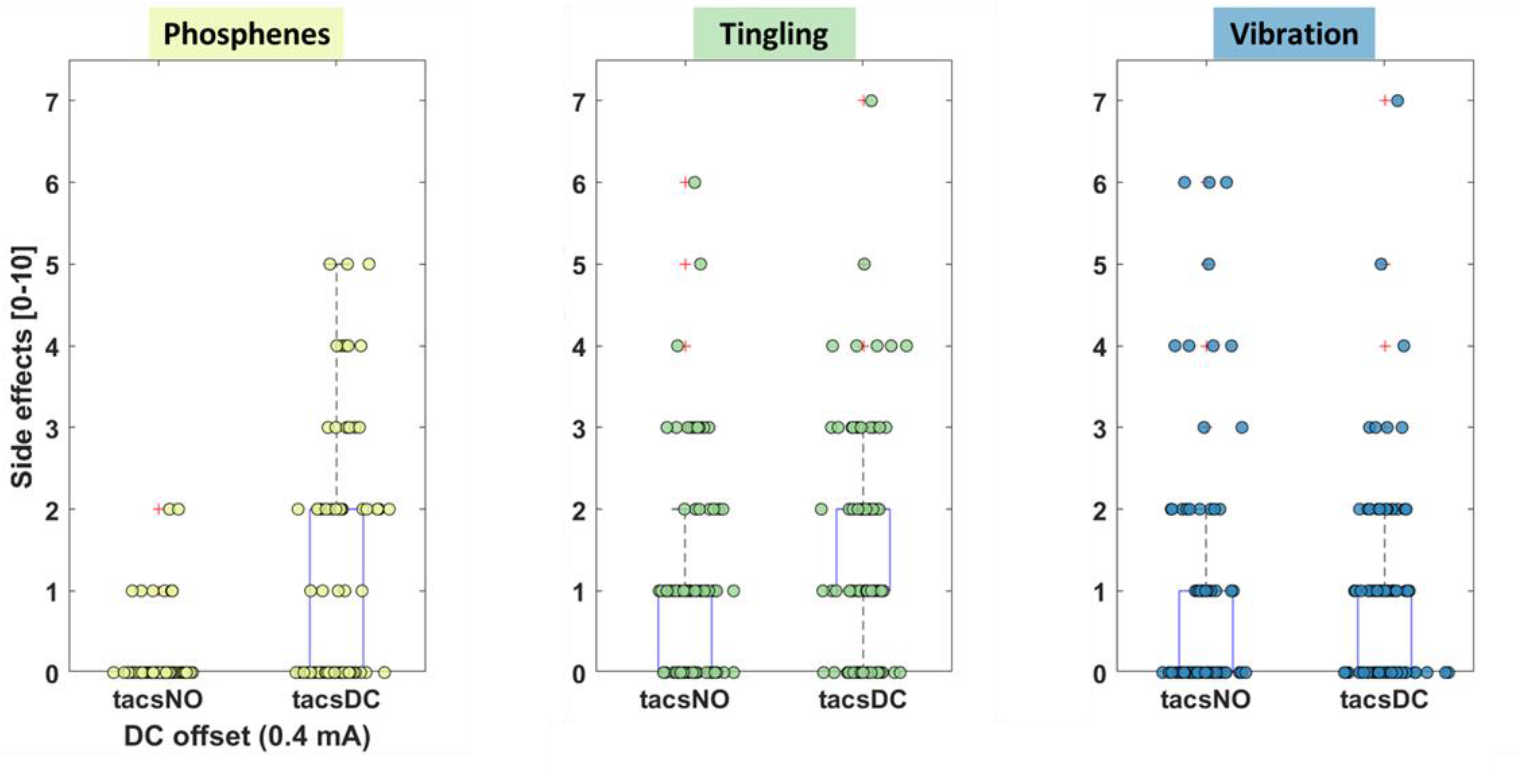
RQ 1.1 Boxplots: Phosphenes, Tingling and Vibration depending on DC-offset. Each point on the boxplots depicts a rating of one participant. Points are jittered for better readability.

### 1.2 SEs differ for at least one frequency

The occurrence of vibration and tingling depended on the stimulation frequency (RQ 1.2). Specifically, compared to the lowest frequency of 250 Hz, higher frequencies were associated with lower levels of both SEs. Phosphenes are slightly reduced in the 250 Hz condition. These results are indicated by a significant main fixed effect for the occurrence of tingling (F(3,214) = 21.42, p <.001) and vibration (F(3,214) = 30.56, p <.001). The analysis of the fixed coefficients revealed that stimulating with 1000 Hz (B=-.724, p=.014), and 2000 Hz (B=-.0669, p=.018) in comparison to 250 Hz (B=1.040, p<.001) led to lower levels of vibration, but not compared to 500 Hz (B=-.395, p=1.119). Tingling values were significantly higher at 250 Hz than at the reference frequency of 2000 Hz (B = 1.196, p < 0.001, Exp(B) = 3.31 95%-KI [2.71, 4.03]). There were no significant differences at 500 Hz, 1000 Hz, and 2000 Hz. Even though there was no significant fixed effect for the occurrence of phosphenes depending on the stimulation frequency, the analysis of the coefficients showed that stimulating with 250 Hz (B=-1.170, p=.006, Exp(B)=.31, 95%-KI [.14-.71] and 500 Hz (B=-1.293, p=.012, Exp(B)=.27, 95%-KI [.10-.75]) in comparison to stimulating with 2000 Hz leads to significantly less phosphenes. Figure 4 provides an overview of the SEs, depending on the stimulation frequency.

**Figure 4:**
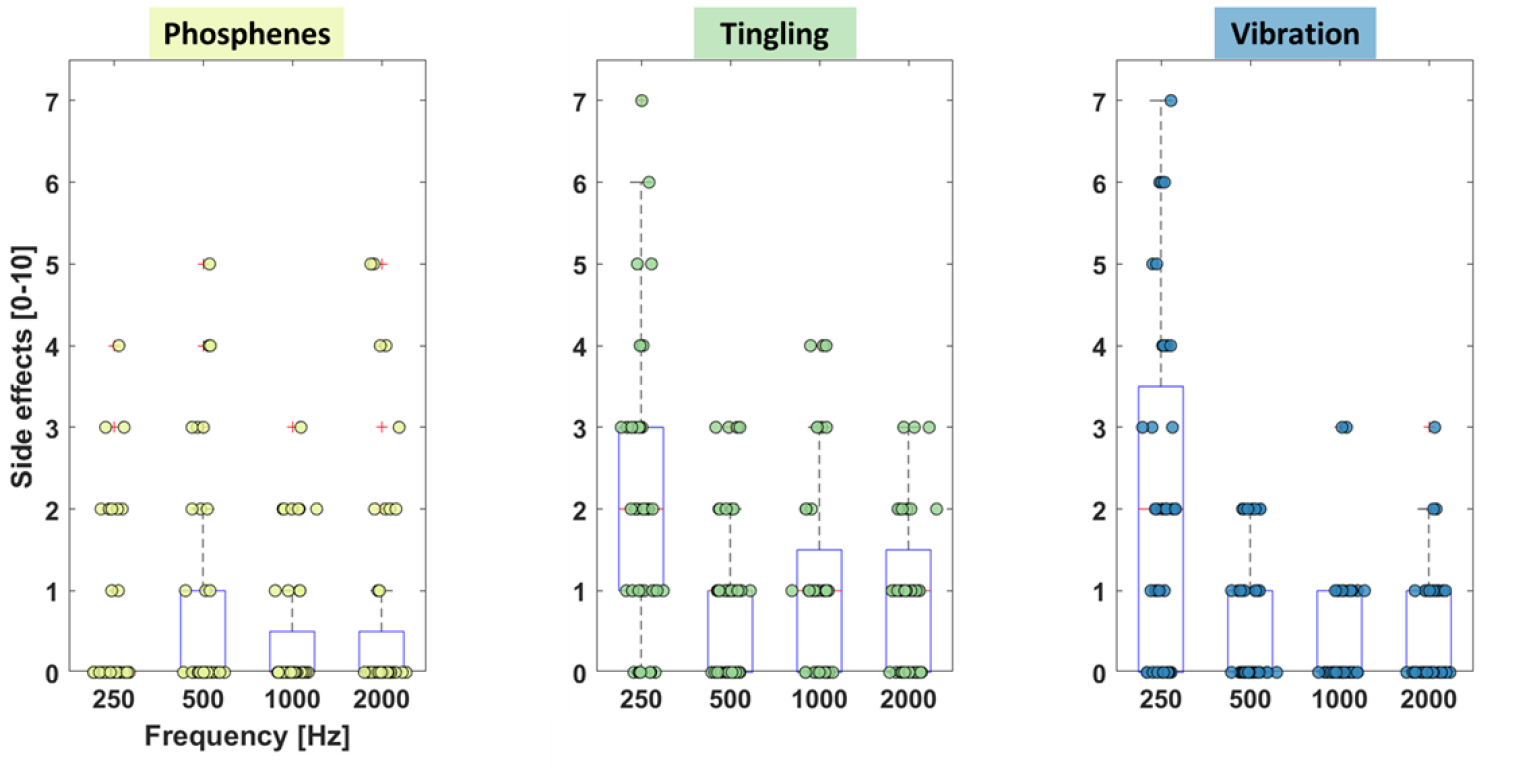
RQ 1.2 Boxplot, Phosphenes, Tingling, Vibration depending on stimulation frequency. Each point on the boxplots depicts a rating of one participant. Points are jittered for better readability.

### 1.3. SEs differ between electrode positions

Sensations of tingling and vibration were slightly increased in the ear-to-ear montage, while phosphenes almost exclusively appeared with a forehead return electrode (RQ1.3).

Those results are indicated by a significant effect for vibration (F(1,214)=12.09, p=.001), tingling (F(1,214)=8.17, p=.005), and phosphenes F(1,214)=15.02, p<.001). The forehead return electrode was associated with significantly lower values for vibration (B=-.511, p<.001, Exp(B)=.6, 95%-KI [.449-.802]) and tingling (B=-.330, p=.005, Exp(B)=.72, 95%-KI [.573-.903]). Phosphenes were primarily present in the forehead montage (B=1.534, p<.001, Exp(B)=4.64, 95%-KI [2.13-10.12]. Figure 5 provides an overview of the SE, depending on return electrode positioning and Figure 6 provides an overview of the medians of the SE scales across stimulation frequencies only and stimulation frequency and DC offset, as well as stimulation frequency and return electrode positions in a heat map.

**Figure 5:**
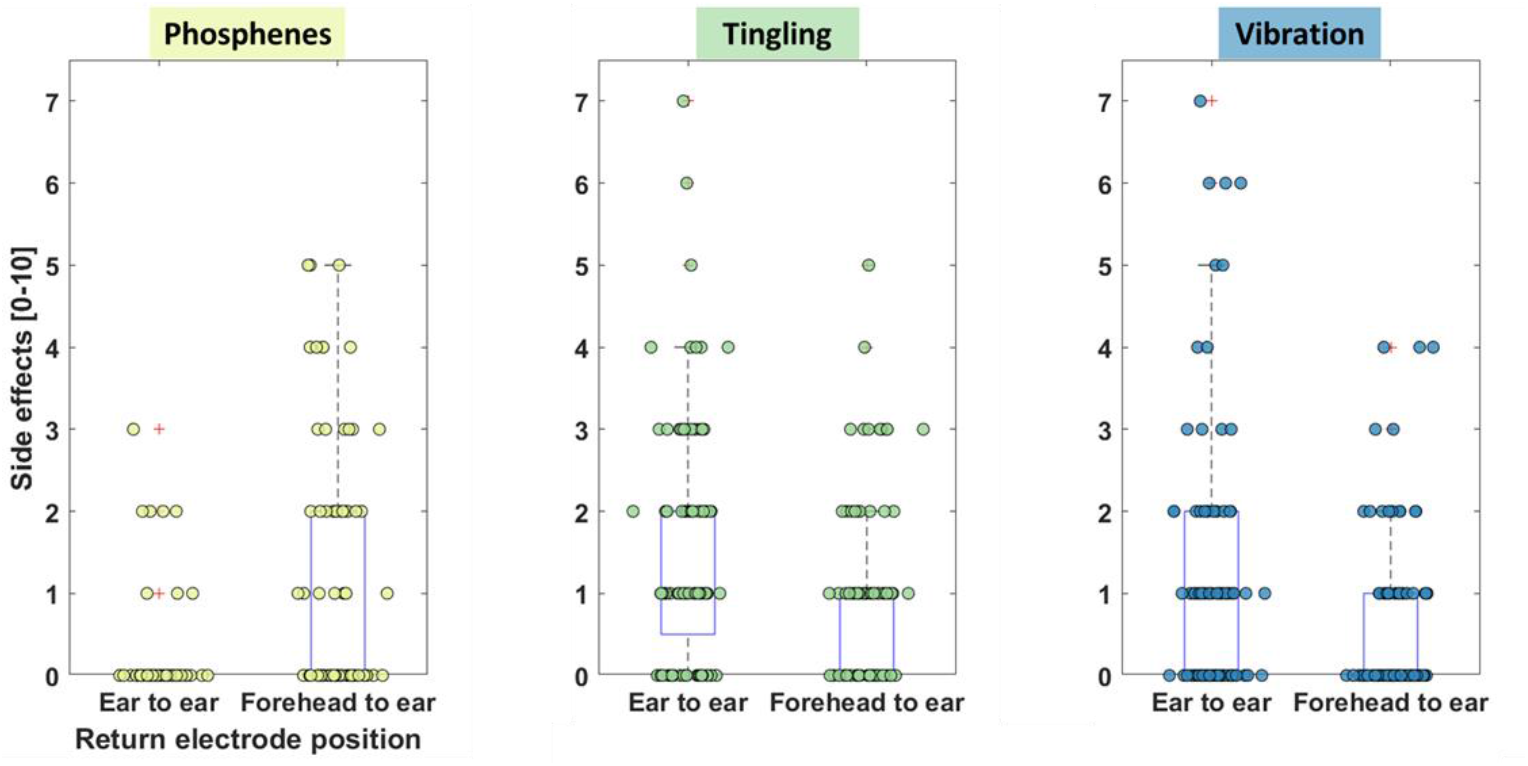
RQ 1.3: Boxplots: Phosphenes, Tingling and Vibration depending on position. Each point on the boxplot represents a rating from one participant. Points are jittered for better readability.

**Figure 6:**
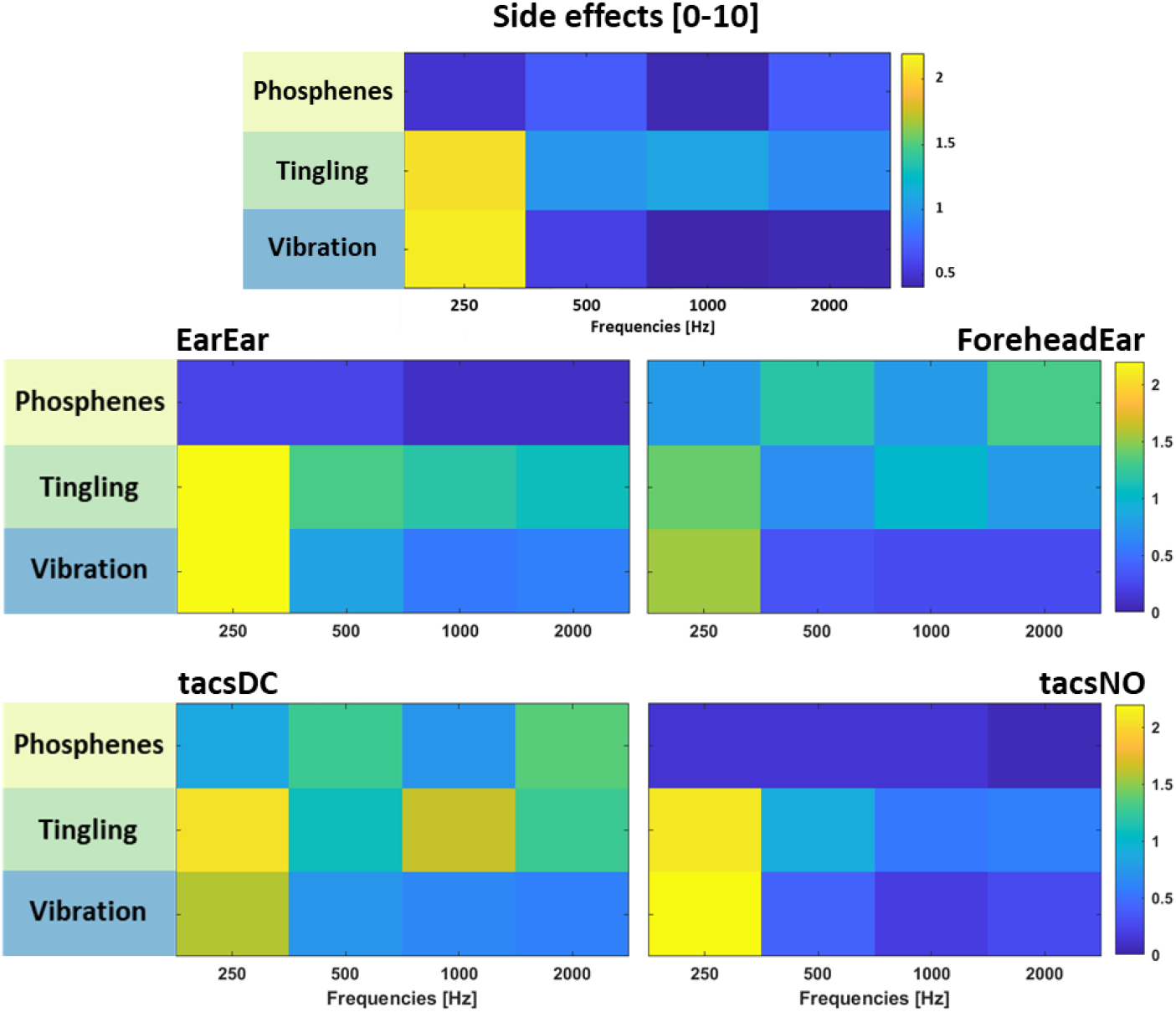
RQ1.1: SE mean across participants depending on stimulation frequency. We present the mean side effect rating across participants (first row), separated by electrode position (RQ 1.2, second row), and by DC offset (RQ 1.3, third row).

Details on the statistical analysis of SEs can be found in the Appendix, Table 1-3.

### 2.1 Hearing impressions do not differ between tacsDC and tacsNO

In terms of DC-offset (RQ 2.1), hearing perception was reported in 85% of the cases with DC-offset compared to 83% without DC-offset (see Figure 7C). There was no significant difference in sound perception between tacsDC and tacsNO (F(1,43)=.010, p =.921, B=.053, p=.921, Exp(B)=1.054, 95%-KI=.363-3.057).

**Figure 7:**
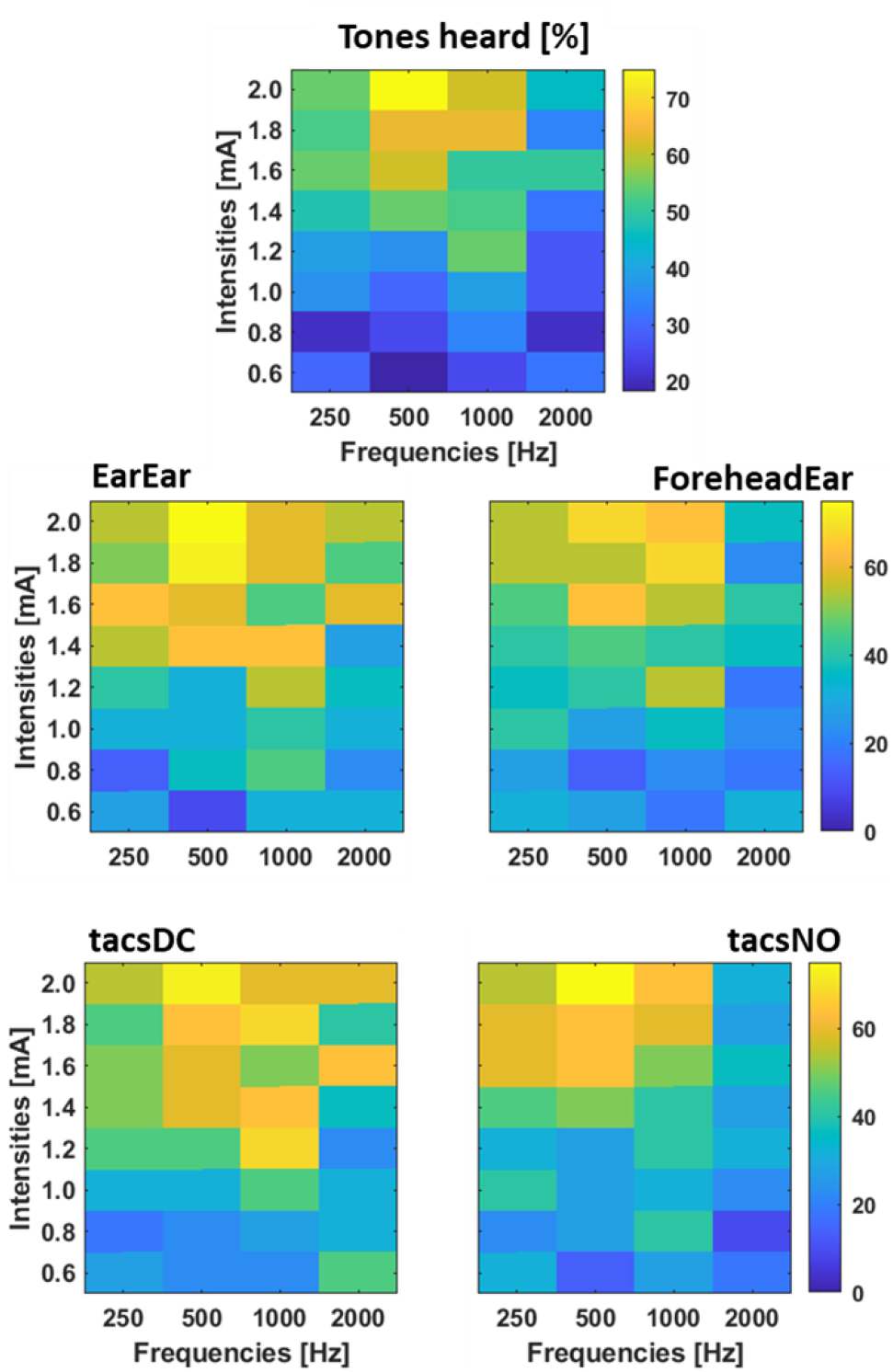
RQ 2: Auditory perception depending on stimulation frequency, electrode position (EarEar=in-ear electrode, ForeheadEar forehead electrode), and DC-offset (tacsDC, tacsNO).

### 2.2 Hearing impressions differ for at least one frequency

Regarding a frequency dependence of hearing perception (RQ 2.2.), no association could be found for fixed effects F(3,43)=.606, p=.615). Nevertheless, it was noticeable that there was a decrease in sound perception in percent when stimulating with higher frequencies (250 Hz=90% > 500 Hz=86% > 1000 Hz= 82% > 2000 Hz= 75% hearing perception) (see Figure 7A). Regarding the fixed coefficients, lower and middle frequencies could be heard more frequently in comparison to higher frequencies (250 Hz: B=1.997, p=.005 Exp(B)= 7.369, 95%-KI [1.867-29.084], 500 Hz: B=2.091, p=.008, Exp(B)= 8.094, 95%-KI [1.796-36.477], 1000 Hz: B=1.590, p=.030, Exp(B)= 4.905, 95%-KI [1.177-20.441], 2000 Hz: B=1.216, p=.069, Exp(B)= 3.372, 95%-KI [0.904-12.574]).

### 2.3. Hearing impressions do not differ between montages

Participants reported hearing the tone more frequently with the return electrode placed in the contralateral ear (90%) than when placed on the forehead (79.1%). In our analysis, we found a tendency towards higher auditory perception with ear-to-ear electrode positioning (see Figure 7B). However, the difference was not significant (F(1,43)=2.535, p=0.119), ear-to-ear: B=(−0.871), p=0.119, Exp(B)=0.419 95%-KI= 0.139-1.261).

Details on the statistical analysis of HIs can be found in the Appendix, Table 4.

### 2.4 Further analysis

#### Distraction

We further investigated whether distraction caused by high SE reduces auditory perception. We did not find a significant link between distraction ratings and HI (F(1,43)=2.761, p=.104, B=1.493, p=.104, Exp(B)=4.450, 95%-KI [0.727-27.241]. We also examined whether the position of the return electrode influenced sound perception. When the return electrode was placed on the forehead, sound was perceived in 52.1% of cases in the left ear, in 23.4% in the right ear, and in 24.5% in both ears. In contrast, with the return electrode positioned in the ear canal, sound perception occurred in 37.4% of cases in the left ear, in 30.3% of cases in the right ear, and in 32.3% of cases in both ears.

#### Dizziness

We quantified the sensation of dizziness by controlling for nystagmus whenever a participant rated dizziness as 1/10 or greater on the Likert-like scale. We did not observe any abnormal eye movements during stimulation that could indicate a disturbance of the vestibular system.

## 3. Discussion

This study shows that in-ear tACS at high frequencies (>80 Hz) targeting peripheral regions of the auditory pathway is well tolerated, consistent with previous findings (Bjekić et al., 2024; Matsumoto & Ugawa, 2017).

Previous studies used electrode setups on the temples, preauricular (contralateral, ipsilateral) on the forehead, and on the top of the head. Regarding SEs, different profiles, depending on the tissue located between the stimulation electrode and the return electrode, have been reported. Among these, ear canal stimulation with a forehead return electrode showed the best efficacy in a single CI patient (Zeng et al., 2019). Despite its possible advantages, no studies have explored ear canal-to-ear canal positioning (Zeng et al., 2011). Our present results show that an ear-to-ear montage slightly increases tingling and vibration, but almost eliminates phosphenes. This setup also reduces preparation time by eliminating the need for hair removal and enabling non-experts to set up home-use devices easily.

We specifically introduced an artificial DC offset and considered its influence.

Second, we examined the SE in relation to stimulation frequency. Facial movements were observed more frequently at the lowest frequency, while tingling and vibration sensations occurred more frequently at lower frequencies overall. Phosphenes tended to occur more frequently in our setup at higher frequencies. This differs from one of the few investigations available in this frequency range: Zeng et al. (2019) found phosphenes to only occur between 5 Hz and 100 Hz. However, both Zeng et al. (2019) and Raco et al. (2014) found phosphenes to occur more likely in higher frequencies within their investigated range, which is consistent with our findings.

Third, we compared the side effect profiles of different return electrodes. Phosphenes were more common with a forehead return electrode, consistent with studies showing visual perception predominantly occurs with frontal or mastoid stimulation, likely due to retinal stimulation (Antal & Paulus, 2013; Schutter & Hortensius, 2010). Symmetrical perceptions are mainly evoked with frontal electrodes, while other electrode positions tend to cause asymmetrical perceptions (Czornik et al., 2022; Raco et al., 2014). Previous studies have primarily described muscle twitches and facial movements resulting from cranial nerve activation (Zeng et al., 2019). Participants rarely reported facial movements in our study. Tingling and dizziness were observed significantly more frequently with ear-to-ear electrode positioning. Dizziness may be due to vestibular stimulation, though dizziness ratings remained low and could not be objectively assessed with the Frenzel goggles.

Regarding hearing perception, we did not find any differences in the intensity threshold required using a DC-offset or stimulating with pure tACS. The findings suggest a potential for both positions to elicit auditory sensations, which does not support the assumption underlying RQ2.1. This stands in contrast to the suggestion by Vanneste et al. (2013) that tACS, unlike tDCS (transcranial direct current stimulation), requires higher stimulation intensities when the auditory cortex is targeted (Vanneste et al., 2013). Regarding RQ 2.2, we found that sound perception gradually decreases at higher frequencies (90% HI at 250 Hz > 75% HI at 2000 Hz). Two mechanisms contribute to this effect. First, tissue acts as a low-pass filter, attenuating higher frequencies while allowing lower frequencies to pass through (Bédard et al., 2006; Gygi & Moschytz, 1997). Second, equal neural responses for different frequencies require equal charge to be delivered. Since charge depends quadratically upon frequency, doubling the frequency of stimulation would require to quadruple the current (Kral, Aplin, & Maier, 2021).

Regarding RQ2.3, we found that placing the return electrode in the ear canal produced more auditory impressions. This finding is consistent with a previous study by Zeng et al., who measured the maximum intensity that reaches the cochlea for ear canal electrodes (Zeng et al., 2019). When the return electrode was placed in the contralateral ear canal, participants reported hearing the sound equally in the right, left, and both ears. This suggests that two ear canal electrodes may stimulate both sides of the auditory system. Speaking to this is the fact that participants were not able to localize the source of the HI when stimulated ear-to-ear.

Discussions about the origin of this HI are still ongoing, since both the cochlea and the auditory nerve might be involved. In other studies, the cochlea was stimulated via electrodes as well as directly via cochlear implants, both of which were effective (Arts et al., 2012; Bovo et al., 2011; Zeng et al., 2011; Zeng et al., 2019). However, the frequency-specificity of the HI makes the cochlea a reasonable target (Zeng et al., 2019).

In summary, our findings demonstrate that in-ear tACS is well-tolerated and capable of eliciting stable auditory perception. The ear-to-ear electrode montage proved to be particularly practical and effective.

### Limitations

Several limitations should be acknowledged. Bilateral stimulation of the auditory system cannot be ruled out in the ear-to-ear montage and the exact side of the stimulation remains uncertain. Using only auditory percepts and SEs as outcomes limits insights into underlying mechanisms. Future studies should address the effectiveness of the approach with a clinical sample.

## 3. Conclusion

In summary, this study found in-ear tACS to reliably produce a stable HI while maintaining a low side effect profile. The SEs caused by DC offset emphasize the need to control for any unintentional influence of Direct Current during tACS. According to our findings, the DC offset is not relevant for evoking hearing impressions in participants, and there are no significant differences in the total stimulation intensity required to evoke hearing impressions. We recommend in-ear return electrode placement as the preferred option. HIs were most frequent at lower frequencies, although it remains to be seen whether the suppression of tinnitus is dependent on matching the stimulation frequency to the tinnitus frequency.

Our study confirms the feasibility of transcranial in-ear stimulation with alternating current. Our findings on favorable stimulation parameters will help guide further research to improve the usability of tACS in tinnitus research. In the future, tinnitus suppression will be analyzed using our preferred setup. By working with patients suffering from chronic tinnitus, further conclusions can be drawn for the practical application of tACS in tinnitus patients.

## Reference list

Antal, A., Boros, K., Poreisz, C., Chaieb, L., Terney, D., & Paulus, W. (2008). Comparatively weak after-effects of transcranial alternating current stimulation (tACS) on cortical excitability in humans. Brain Stimulation, 1(2), 97–105. 10.1016/j.brs.2007.10.001

Antal, A., & Paulus, W. (2013). Transcranial alternating current stimulation (tACS) [Review]. Frontiers in Human Neuroscience, Volume 7 - 2013. 10.3389/fnhum.2013.00317

Arshad, Q., Nigmatullina, Y., Roberts, R. E., Bhrugubanda, V., Asavarut, P., & Bronstein, A. M. (2014). Left cathodal trans-cranial direct current stimulation of the parietal cortex leads to an asymmetrical modulation of the vestibular-ocular reflex. Brain Stimulation, 7(1), 85–91. 10.1016/j.brs.2013.07.002

Arts, R. A., George, E. L., Janssen, M., Griessner, A., Zierhofer, C., & Stokroos, R. J. (2016). Tinnitus Suppression by Intracochlear Electrical Stimulation in Single Sided Deafness - A Prospective Clinical Trial: Follow-Up. PLoS One, 11(4), e0153131. 10.1371/journal.pone.0153131

Arts, R. A. G. J., George, E. L. J., Stokroos, R. J., & Vermeire, K. (2012). Review: cochlear implants as a treatment of tinnitus in single-sided deafness. Current Opinion in Otolaryngology & Head and Neck Surgery, 20(5). 10.1097/MOO.0b013e3283577b66

Auerbach, B. D., Rodrigues, P. V., & Salvi, R. J. (2014). Central gain control in tinnitus and hyperacusis. Frontiers in Neurology, 5, 206. 10.3389/fneur.2014.00206

Bédard, C., Kröger, H., & Destexhe, A. (2006). Model of low-pass filtering of local field potentials in brain tissue. Physical Review E, 73(5 Pt 1), 051911. 10.1103/PhysRevE.73.051911

Biswas, R., Lugo, A., Akeroyd, M. A., Schlee, W., Gallus, S., & Hall, D. A. (2022). Tinnitus prevalence in Europe: a multi-country cross-sectional population study. The Lancet Regional Health Europe, 12, 100250. 10.1016/j.lanepe.2021.100250

Bjekić, J., Živanović, M., Stanković, M., Paunović, D., Konstantinović, U., & Filipović, S. R. (2024). The subjective experience of transcranial electrical stimulation: a within-subject comparison of tolerability and side effects between tDCS, tACS, and otDCS [Brief Research Report]. Frontiers in Human Neuroscience, Volume 18 - 2024. 10.3389/fnhum.2024.1468538

Bovo, R., Ciorba, A., & Martini, A. (2011). Tinnitus and cochlear implants. Auris Nasus Larynx, 38(1), 14–20. 10.1016/j.anl.2010.05.003

Chaieb, L., Antal, A., Pisoni, A., Saiote, C., Opitz, A., Ambrus, G. G., Focke, N., & Paulus, W. (2014). Safety of 5 kHz tACS. Brain Stimulation, 7(1), 92–96. 10.1016/j.brs.2013.08.004

Chang, J. E., & Zeng, F. G. (2012). Tinnitus suppression by electric stimulation of the auditory nerve. Front Syst Neurosci, 6, 19. 10.3389/fnsys.2012.00019

Corp., I. (Released 2023). IBM SPSS Statistics for Macintosh. In (Version Version 29.0.2.0) IBM Corp.

Czornik, M., Malekshahi, A., Mahmoud, W., Wolpert, S., & Birbaumer, N. (2022). Psychophysiological treatment of chronic tinnitus: A review. Clinical Psychology & Psychotheraphy, 29(4), 1236–1253. 10.1002/cpp.2708

Dauman, R., & Bouscau-Faure, F. (2005). Assessment and amelioration of hyperacusis in tinnitus patients. Acta Oto-Laryngologica, 125(5), 503–509. 10.1080/00016480510027565

Elyamany, O., Leicht, G., Herrmann, C. S., & Mulert, C. (2021). Transcranial alternating current stimulation (tACS): from basic mechanisms towards first applications in psychiatry. European Archives of Psychiatry and Clinical Neuroscience, 271(1), 135–156. 10.1007/s00406-020-01209-9

Gygi, A. E., & Moschytz, G. S. (1997). Low-pass filter effect in the measurement of surface EMG. In Proceedings of Computer Based Medical Systems (pp. 183–188). IEEE.

Hackenberg, B., O’Brien, K., Döge, J., Lackner, K. J., Beutel, M. E., Münzel, T., Pfeiffer, N., Schulz, A., Schmidtmann, I., Wild, P. S., Matthias, C., & Bahr-Hamm, K. (2023). Tinnitus Prevalence in the Adult Population—Results from the Gutenberg Health Study. Medicina, 59. 10.3390/medicina59030620

Heller, A. J. (2003). Classification and epidemiology of tinnitus. Otolaryngologic Clinics of North America, 36(2), 239–248. 10.1016/S0030-6665(02)00160-3

Holder, J. T., O’Connell, B., Hedley-Williams, A., & Wanna, G. (2017). Cochlear implantation for single-sided deafness and tinnitus suppression. American Journal of Otolaryngology, 38(2), 226–229. 10.1016/j.amjoto.2017.01.020

Jarach, C. M., Lugo, A., Scala, M., van den Brandt, P. A., Cederroth, C. R., Odone, A., Garavello, W., Schlee, W., Langguth, B., & Gallus, S. (2022). Global Prevalence and Incidence of Tinnitus: A Systematic Review and Meta-analysis. JAMA Neurology, 79(9), 888–900. 10.1001/jamaneurol.2022.2189

Jastreboff, P. J., & Jastreboff, M. M. (2003). Tinnitus Retraining Therapy for patients with tinnitus and decreased sound tolerance. Otolaryngologic Clinics of North America, 36(2), 321–336. 10.1016/S0030-6665(02)00172-X

Kaltenbach, J. A., Zhang, J., & Finlayson, P. (2005). Tinnitus as a plastic phenomenon and its possible neural underpinnings in the dorsal cochlear nucleus. Hearing Research, 206(1), 200–226. 10.1016/j.heares.2005.02.013

Kral, A., Aplin, F., & Maier, H. (2021). Prostheses for the brain: introduction to neuroprosthetics. Academic Press.

Laakso, I., Nissi, J., & Kangasmaa, O. (2025). Vestibular involvement in transcranial electrical stimulation: Body sway as a marker of unintended stimulation. Brain Stimulation, 18(1), 34–36. 10.1016/j.brs.2024.12.1188

Landgren, S., & Cajander, Å. (2021). Non-use of Digital Health Consultations Among Swedish Elderly Living in the Countryside [Original Research]. Frontiers in Public Health, 9. 10.3389/fpubh.2021.588583

Langguth, B., & De Ridder, D. (2013). Tinnitus: therapeutic use of superficial brain stimulation. Handbook of Clinical Neurology, 116, 441–467. 10.1016/b978-0-444-53497-2.00036-x

Matsumoto, H., & Ugawa, Y. (2017). Adverse events of tDCS and tACS: A review. Clinical Neurophysiology Practice, 2, 19–25. 10.1016/j.cnp.2016.12.003

Mazurek, B., Hesse, G., Dobel, C., Kratzsch, V., Lahmann, C., & Sattel, H. (2022). Chronic Tinnitus: Diagnosis and Treatment. Deutsches Ärzteblatt International, 119(13), 219–225. doi: 10.3238/arztebl.m2022.0135

McCormack, A., Edmondson-Jones, M., Somerset, S., & Hall, D. (2016). A systematic review of the reporting of tinnitus prevalence and severity. Hearing Research, 337, 70–79. 10.1016/j.heares.2016.05.009

Neff, P., Michels, J., Meyer, M., Schecklmann, M., Langguth, B., & Schlee, W. (2017). 10 Hz Amplitude Modulated Sounds Induce Short-Term Tinnitus Suppression. Frontiers in Aging Neuroscience, 9, 130. 10.3389/fnagi.2017.00130

Noreña, A. J. (2011). An integrative model of tinnitus based on a central gain controlling neural sensitivity. Neuroscience & Biobehavioral Reviews, 35(5), 1089–1109. 10.1016/j.neubiorev.2010.11.003

Park, K. W., Kullar, P., Malhotra, C., & Stankovic, K. M. (2023). Current and Emerging Therapies for Chronic Subjective Tinnitus. Journal of Clinical Medicine, 12(20). 10.3390/jcm12206555

Quaranta, N., Fernandez-Vega, S., D’Elia, C., Filipo, R., & Quaranta, A. (2008). The effect of unilateral multichannel cochlear implant on bilaterally perceived tinnitus. Acta Otolaryngol, 128(2), 159–163. 10.1080/00016480701387173

Raco, V., Bauer, R., Olenik, M., Brkic, D., & Gharabaghi, A. (2014). Neurosensory effects of transcranial alternating current stimulation. Brain Stimulation, 7(6), 823–831. 10.1016/j.brs.2014.08.005

Ruckenstein, M. J., Hedgepeth, C., Rafter, K. O., Montes, M. L., & Bigelow, D. C. (2001). Tinnitus suppression in patients with cochlear implants. Otology & Neurotology, 22(2), 200–204. 10.1097/00129492-200103000-00014

Schutter, D. J. L. G., & Hortensius, R. (2010). Retinal origin of phosphenes to transcranial alternating current stimulation. Clinical Neurophysiology, 121(7), 1080–1084. 10.1016/j.clinph.2009.10.038

Suh, M.-W., Tran, P., Richardson, M., Sun, S., Xu, Y., Djalilian, H. R., Lin, H. W., & Zeng, F.-G. (2022). Electric hearing and tinnitus suppression by noninvasive ear stimulation. Hearing Research, 415, 108431. 10.1016/j.heares.2022.108431

Vanneste, S., Walsh, V., Van De Heyning, P., & De Ridder, D. (2013). Comparing immediate transient tinnitus suppression using tACS and tDCS: a placebo-controlled study. Experimental Brain Research, 226(1), 25–31. 10.1007/s00221-013-3406-7

Zeng, F.-G. (2013). An active loudness model suggesting tinnitus as increased central noise and hyperacusis as increased nonlinear gain. Hearing Research, 295, 172–179. 10.1016/j.heares.2012.05.009

Zeng, F.-G., Tang, Q., Dimitrijevic, A., Starr, A., Larky, J., & Blevins, N. H. (2011). Tinnitus suppression by low-rate electric stimulation and its electrophysiological mechanisms. Hearing Research, 277(1), 61–66. 10.1016/j.heares.2011.03.010

Zeng, F. G., Richardson, M., Tran, P., Lin, H., & Djalilian, H. (2019). Tinnitus Treatment Using Noninvasive and Minimally Invasive Electric Stimulation: Experimental Design and Feasibility. Trends in Hearing, 23, 2331216518821449. 10.1177/2331216518821449

Zeng, F. G., Tran, P., Richardson, M., Sun, S., & Xu, Y. (2019). Human Sensation of Transcranial Electric Stimulation. Scientific Reports, 9(1), 15247. 10.1038/s41598-019-51792-8

