## Appendix for "In-ear-tACS: Auditory Perception and Side Effects depend on Electrode Montage, Frequency, and DC-Offset"

*Table 1: GLMM Fixed Effects Table for vibration (reference position: 2, reference DC-offset (tacsNO))*

| *Fixed Effect* | *B (Coefficient)*[1] | *SE*[2] | *t*[3] | *p-value*[4] | *95% CI (B)*[5] | *Exp (B) (OR)*[6] | *95% CI (OR)*[7] |
|---|---|---|---|---|---|---|---|
| freq[8]= 250 Hz | 1.04 | .123 | 8.437 | 0.0 | (-.797-.283) | 2.829 | (2.219-3.608) |
| freq[8]= 500 Hz | -.395 | .252 | -1.565 | 0.119 | (-.892-.102) | .674 | (.41-1.108) |
| freq[8]=1000 Hz | -.724 | .291 | -2.48 | .014 | (-1.299-(.149)) | .485 | (.273-.862) |
| freq[8]= 2000 Hz | -.669 | .285 | -2.347 | .019 | (-1.231-(-.108)) | .512 | (.292-.897) |
| pos[9]=1 | -.511 | .146 | -3.477 | .001 | (-.8-(-.221)) | .6 | (.449-.802) |
| DC[10]=1 | .103 | .142 | .723 | .47 | (-.178-.384) | 1.109 | (.837-1.468) |

*B (Coefficient) = unstandardized regression coefficient*[1]
*SE = standard error*[2]
*t = test statistic (t-value)*[3]
*p-value = significance level*[4]
*95%-CI (B) = 95% confidence interval for B*[5]
*Exp(B) (OR) = exponentiated coefficient, representing the odds ratio (OR)*[6]
*95% CI (OR) = 95% confidence interval for the odds ratio*[7]
*freq= stimulation frequency (Hz)*[8]
*pos = return electrode position (1= forehead, 2=ear canal)*[9]
*DC = direct current offset condition (0=tacsNO, 1=tacsDC)*[10]

*Table 2: GLMM Fixed Effects Table for tingling (reference position: 2, reference DC-offset (tacsNO))*

| *Fixed Effect* | *B (Coefficient)*[1] | *SE*[2] | *t*[3] | *p-value*[4] | *95% CI (B)*[5] | *Exp (B) (OR)*[6] | *95% CI (OR)*[7] |
|---|---|---|---|---|---|---|---|
| freq[8]= 250 Hz | 1.196 | 0.1006 | 11.883 | 0.0 | (0.998-1.394) | 3.307 | (2.712-4.032) |
| freq[8]= 500 Hz | 0.244 | 0.177 | 1.372 | 0.171 | (-0.106-0.594) | 1.276 | (0.899-1.812) |
| freq[8]=1000 Hz | 0.331 | 0.171 | 1.931 | 0.055 | (-0.0007-0.669) | 1.392 | (0.993-1.952) |
| freq[8]= 2000 Hz | 0.173 | 0.183 | 0.946 | 0.345 | (-0.188-0.534) | 1.189 | (0.829-1.706) |
| pos[9]=1 | -0.33 | 0.115 | -2.858 | 0.005 | (-0.557-(-0.102) | 0.719 | (0.573-0.903) |
| DC[10]=1 | -0.195 | 0.114 | -1.701 | 0.09 | (-0.42-0.031) | 0.823 | (0.657-1.031) |

*B (Coefficient) = unstandardized regression coefficient*[1]
*SE = standard error*[2]
*t = test statistic (t-value)*[3]
*p-value = significance level*[4]
*95%-CI (B) = 95% confidence interval for B*[5]
*Exp(B) (OR) = exponentiated coefficient, representing the odds ratio (OR)*[6]
*95% CI (OR) = 95% confidence interval for the odds ratio*[7]
*freq= stimulation frequency (Hz)*[8]
*pos = return electrode position (1= forehead, 2=ear canal)*[9]
*DC = direct current offset condition (0=tacsNO, 1=tacsDC)*[10]

*Table 3: GLMM Fixed Effects Table for phosphenes (reference position: 2, reference DC-offset (tacsNO))*

| *Fixed Effect* | *B (Coefficient)[1]* | *SE[2]* | *t[3]* | *p-value[4]* | *95% CI (B)[5]* | *Exp (B) (OR)[6]* | *95% CI (OR)[7]* |
|---|---|---|---|---|---|---|---|
| freq[8]= 250 Hz | -1.17 | 0.418 | -2.8 | 0.006 | (-1.993-(-0.346)) | 0.31 | (0.136-0.707) |
| freq[8]= 500 Hz | -0.804 | 0.447 | -1.197 | 0.074 | (-1.686-0.078) | 0.448 | (0.185-1.081 |
| freq[8]=1000 Hz | -1.293 | 0.508 | -2.547 | 0.012 | (-2.294-(-0.293)) | 0.274 | (0.101-0.746) |
| freq[8]= 2000 Hz | -0.804 | 0.447 | -1.797 | 0.074 | (-1.686-0.078) | 0.448 | (0.185-1.081) |
| pos[9]=1 | 1.534 | 0.396 | -3.876 | 0.0 | (0.754-2.314) | 4.636 | (2.125-10.115) |
| DC[10]=1 | -2.145 | 0.493 | -4.345 | 0.0 | (-3.117-(-1.172)) | 0.117 | (0.044-0.31) |

*B (Coefficient) = unstandardized regression coefficient[1]*
*SE = standard error[2]*
*t = test statistic (t-value)[3]*
*p-value = significance level[4]*
*95%-CI (B) = 95% confidence interval for B[5]*
*Exp(B) (OR) = exponentiated coefficient, representing the odds ratio (OR)[6]*
*95% CI (OR) = 95% confidence interval for the odds ratio[7]*
*freq= stimulation frequency (Hz)[8]*
*pos = return electrode position (1= forehead, 2=ear canal)[9]*
*DC = direct current offset condition (0=tacsNO, 1=tacsDC)[10]*

*Table 4: GLMM Fixed Effects for auditory perception (DC-offset reference tacsDC, stimulation frequency reference 2000 Hz)*

| *Fixed Effect* | *B (Coefficient)[1]* | *SE[2]* | *t[3]* | *p-value[4]* | *95% CI (B)[5]* | *Exp (B) (OR)[6]* | *95% CI (OR)[7]* |
|---|---|---|---|---|---|---|---|
| freq[8]= 250 Hz | -1.831 | 0.692 | -2.646 | 0.009 | (-3.196-(-0.466)) | 0.16 | (0.041-0.627) |
| freq[8]= 500 Hz | -1.434 | 0.756 | -1897 | 0.059 | (-2.925-0.057) | 0.238 | (0.054-1.058) |
| freq[8]=1000 Hz | -0.787 | 0.721 | -1.092 | 0.276 | (-2.208-0.635) | 0.455 | (0.11-1.887) |
| freq[8]= 2000 Hz | 0.0 (Ref) | | | | | 1.0 | |
| pos[9]=1 | -0.724 | 0.92 | -0.788 | 0.432 | (-2.538-1.089) | 0.485 | (0.079-2.972) |
| pos[9]=2 | -2.074 | 0.957 | -2.167 | 0.031 | (-3.961-(-0.186) | 0.126 | (0.019-0.83) |
| DC[10]=0 | 0.087 | 0.503 | 0.173 | 0.863 | (-0.904-1.078) | 1.091 | (0.405-2.939 |
| distraction | 0.83 | 0.762 | -1.089 | 0.278 | (-2.332-0.673) | 0.436 | (0.097-1.96) |

*B (Coefficient) = unstandardized regression coefficient[1]*
*SE = standard error[2]*
*t = test statistic (t-value)[3]*
*p-value = significance level[4]*
*95%-CI (B) = 95% confidence interval for B[5]*
*Exp(B) (OR) = exponentiated coefficient, representing the odds ratio (OR)[6]*
*95% CI (OR) = 95% confidence interval for the odds ratio[7]*
*freq= stimulation frequency (Hz)[8]*
*pos = return electrode position (1= forehead, 2=ear canal)[9]*
*DC = direct current offset condition (0=tacsNO, 1=tacsDC)[10]*
